# Reclassification of *Brucella ciceri* as later heterotypic synonyms of *Brucella intermedia*

**DOI:** 10.1101/2020.08.16.251660

**Authors:** Ayixon Sánchez-Reyes

## Abstract

Recently Hördt et al. 2020 proposed to merge *Ochrobactrum* and *Brucella* genera based on up to date phylogenomic evidence and overall genomic divergence among *Brucella-Ochrobactrum* clade. This led to the description of the new combinations *Brucella ciceri* comb. nov., basonym: *Ochrobactrum ciceri* Imran et al. 2010 and *Brucella intermedia* comb. nov., basonym: *Ochrobactrum intermedium* Velasco et al. 1998. However, the type species for *Brucella ciceri* DSM 22292^T^ and *Brucella intermedia* LMG 3301^T^ show whole-genome coherence at the species level (ANI = 98.21 %, Mash D = 0.0154006, dDDH relatedness >70%), suggesting that may belong to the same genomospecies. Also, both taxa formed a single clade in the phylogenomic tree based on single-copy gene sequences. Previously reported phenotypic data offer a context where both taxa are highly related supporting this synonymy. Therefore, *Brucella ciceri* should be reclassified as later heterotypic synonyms of *Brucella intermedia*, which has priority. The species description is consequently amended.

## Introduction

*Brucella* genus Meyer and Shaw 1920 (Approved Lists 1980) (1) is a member of the α-Proteobacteria class, with several species historically recognized as virulent for humans. At present, possess 25 child taxa with validly published names and a least five synonyms. Recently, Hördt et al. (2) analyzed more 1000 α-Proteobacteria type-strain genome composites, and suggested major emendation’s for many species according to phylogenetic features, resting on genomic data. One of the changes suggested by these authors includes the genus *Ochrobactrum* Holmes et al. 1988 (3) as part of the *Brucella* group, as well as type species of *Ochrobactrum* seems to be a clade more closely related to *Brucella*. These changes were validated in the announcement of new names and new combinations published by Oren and Garrity. (4), so, *Ochrobactrum* actually constitutes a synonym of *Brucella*. This led to the description of the new combinations *Brucella ciceri* comb. nov., basonym: *Ochrobactrum ciceri* Imran et al. 2010 (5) and *Brucella intermedia* comb. nov., basonym: *Ochrobactrum intermedium* Velasco et al. 1998 (6).

However, during a comparative analysis of 30 environmental genomes of the *Brucellaceae* family (unpublished data), arose that the type species for *B. ciceri* DSM 22292^T^ and *B. intermedia* LMG 3301^T^ share whole-genome coherence at the species level (ANI = 98.21 %, Mash D = 0.0154006, dDDH relatedness >70%), suggesting that may belong to the same genomospecies. Both taxa formed a single clade in the phylogenomic tree based on single-copy gene sequences and form a single species-specific cluster according to several genomic properties. In an independent study, Ashford et al. (7) also found high levels of similarity between type strains of synonym’s *O. ciceri* and *O. intermedium* and cautiously suggest a possible re-classification as a single species. Therefore, this note formally proposes that *Brucella ciceri* DSM 22292^T^ should be reclassified as later heterotypic synonyms of *Brucella intermedia* LMG 3301^T^ which has priority and present an emended species description.

## Materials and Methods

### Determination of closely related type strains

The *B. ciceri* DSM 22292^T^ query genome was compared against all type strain genomes accessible in the Type (Strain) Genome Server (TYGS) (https://tygs.dsmz.de) (8), through the MASH tool, which estimate the mutational rate between representative MinHash sketches (9). The ten type strains hits with the lowest MASH distances were chosen for further analysis using the Genome BLAST Distance Phylogeny approach (GBDP) under the algorithm ‘coverage’ and distance formula d5 (10).

### Pairwise comparison of genome sequences

All pairwise genome comparisons were executed using GBDP distance functions. Digital DDH values and confidence intervals were calculated using the recommended settings of the GGDC 2.1 (10). The type-based species clustering was performed employing a 70% dDDH radius among strains with standing in nomenclature. Average nucleotide identity was calculated with fastANI algorithm (11).

### Phylogenetic inference

A minimum evolution tree was inferred via FASTME 2.1.4 according to (12). Branch support values were inferred with 100 pseudo-bootstrap replicates each. The trees were rooted at the midpoint.

## Results and discussion

Whole-genome similarity metrics help to solve taxonomic issues, through the comparison of conserved sequence elements among different genomes. In this work, I have applied several alignment-free sequence mapping tools, to estimate the correct taxonomic status for the strain *B. ciceri* DSM 22292^T^ (Alphaproteobacteria; Rhizobiales; Brucellaceae). Contrary to what is established, *B. ciceri* DSM 22292^T^ does not appear to be a separate species within the *Brucellaceae* family; it shares ~0.015 of Mash distance and ~98.2% of ANI with all *B. intermedia* types genomes, confirming a clear intra-species boundaries prevalence between these two strains (Table 1). As expected, Mash distance ≤ 0.05 correlates well with ANI, and both metrics support species-specific delineation in prokaryotes. These results also reveal a clear genetic gap among all *B. intermedia* types genomes and the rest of the genomes analyzed for the genus, which allows clustering according to genome similarity (9,11).

**Table 1.**
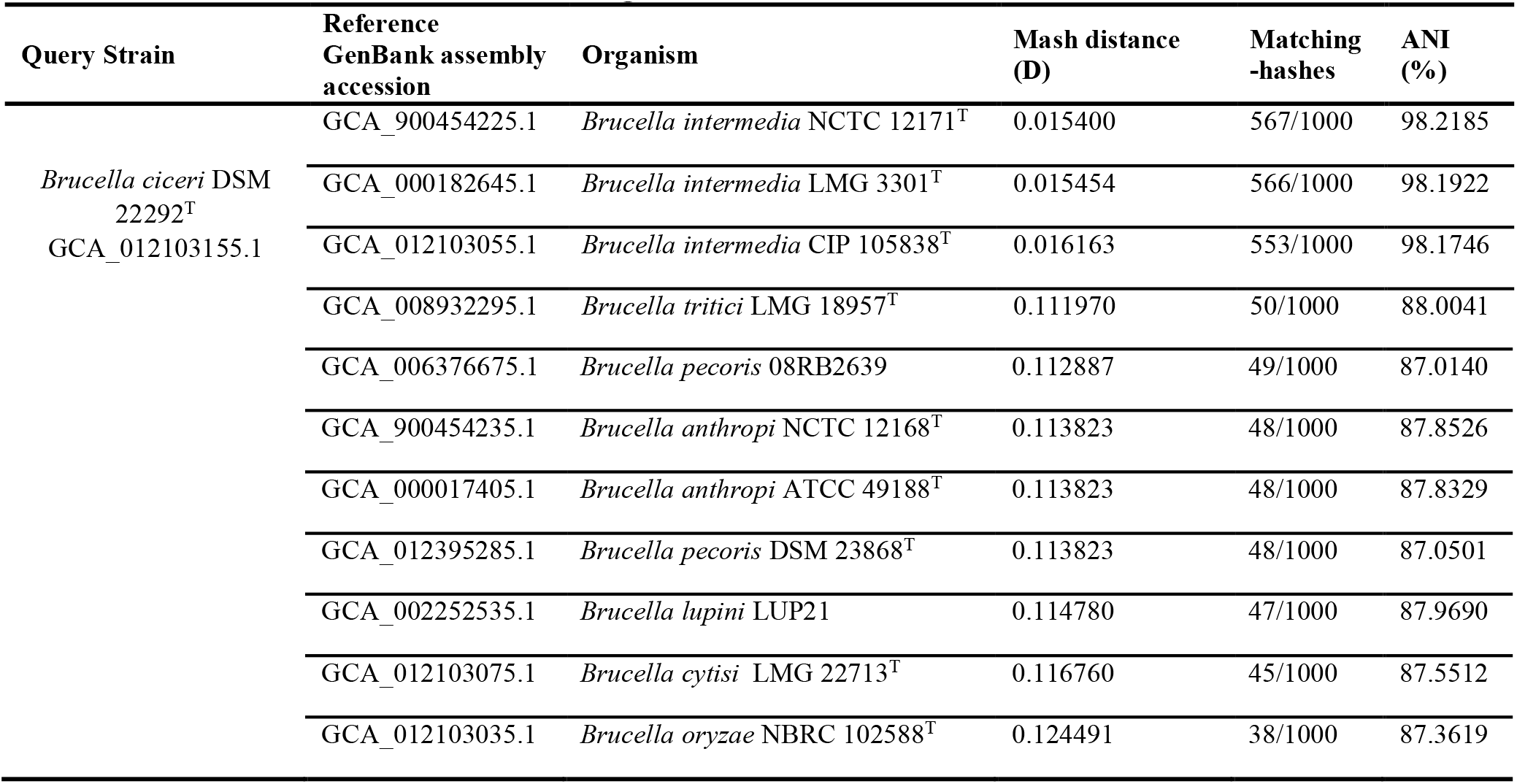
Genomic distance estimation using MinHash

The estimation of DNA-DNA hybridization *in-silico* offers a compelling indication of the species-specific relationship between *B. ciceri* and *B. intermedia* (Table 2). The dDDH (genome-to-genome) comparison shows that the strain DSM 22292^T^ was assigned to the *B. intermedia* species cluster; with all dDDH metrics suggesting a reliable demarcation (CI and dDDH > 80%). Also, both strains share minimal G+C difference supporting they belong to the same species (10).

**Table 2:**
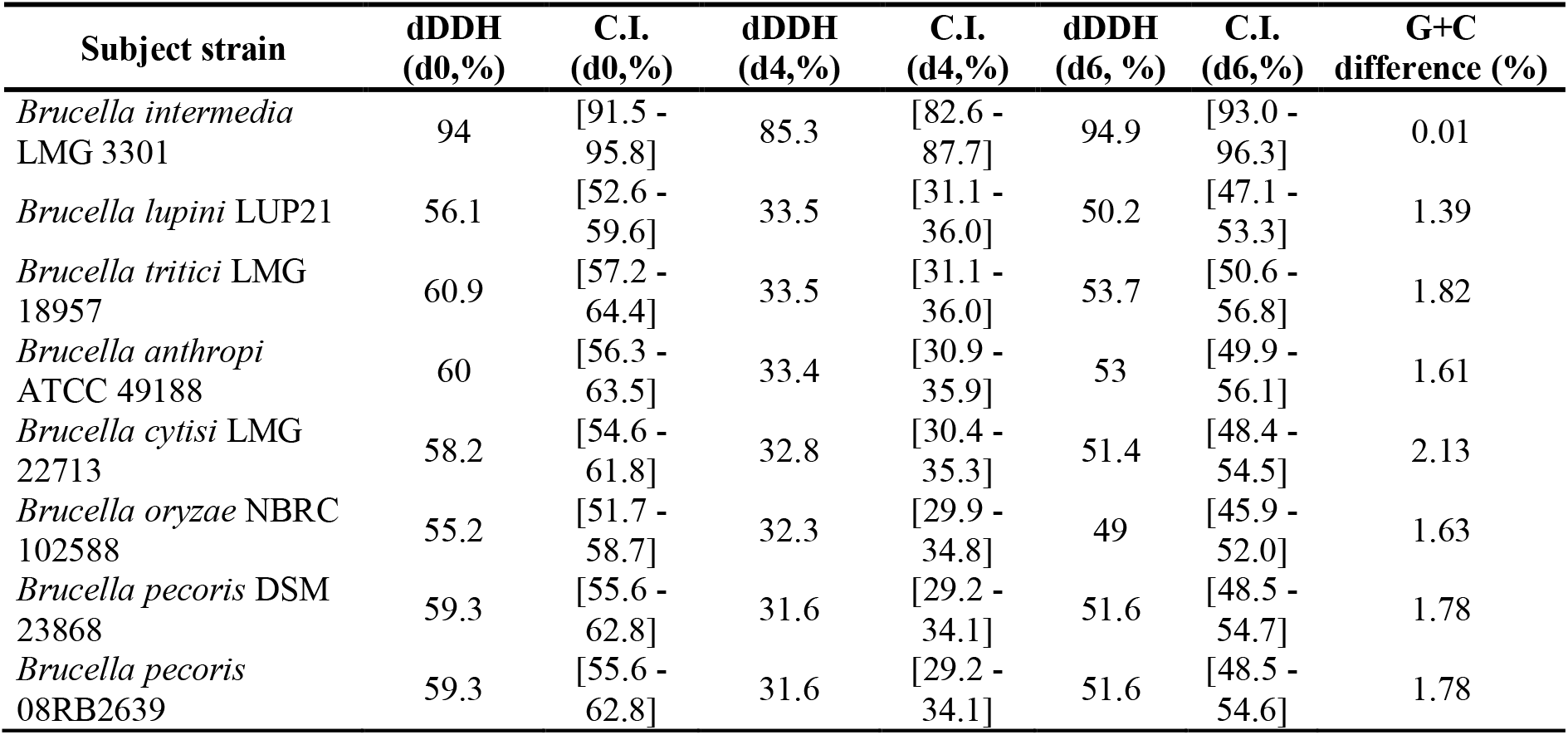
Pairwise comparisons of *Brucella ciceri* DSM 22292^T^ genome vs. related type strain genomes. The table is ordered by column dDDH (d4) in descending order

| Subject strain | dDDH<br>(d0,%) | C.I.<br>(d0,%) | dDDH<br>(d4,%) | C.I.<br>(d4,%) | dDDH<br>(d6, %) | C.I.<br>(d6,%) | G+C<br>difference (%) |
| --- | --- | --- | --- | --- | --- | --- | --- |
| *Brucella intermedia*<br>LMG 3301 | 94 | [91.5 -<br>95.8] | 85.3 | [82.6 -<br>87.7] | 94.9 | [93.0 -<br>96.3] | 0.01 |
| *Brucella lupini* LUP21 | 56.1 | [52.6 -<br>59.6] | 33.5 | [31.1 -<br>36.0] | 50.2 | [47.1 -<br>53.3] | 1.39 |
| *Brucella tritici* LMG<br>18957 | 60.9 | [57.2 -<br>64.4] | 33.5 | [31.1 -<br>36.0] | 53.7 | [50.6 -<br>56.8] | 1.82 |
| *Brucella anthropi*<br>ATCC 49188 | 60 | [56.3 -<br>63.5] | 33.4 | [30.9 -<br>35.9] | 53 | [49.9 -<br>56.1] | 1.61 |
| *Brucella cytisi* LMG<br>22713 | 58.2 | [54.6 -<br>61.8] | 32.8 | [30.4 -<br>35.3] | 51.4 | [48.4 -<br>54.5] | 2.13 |
| *Brucella oryzae* NBRC<br>102588 | 55.2 | [51.7 -<br>58.7] | 32.3 | [29.9 -<br>34.8] | 49 | [45.9 -<br>52.0] | 1.63 |
| *Brucella pecoris* DSM<br>23868 | 59.3 | [55.6 -<br>62.8] | 31.6 | [29.2 -<br>34.1] | 51.6 | [48.5 -<br>54.7] | 1.78 |
| *Brucella pecoris*<br>08RB2639 | 59.3 | [55.6 -<br>62.8] | 31.6 | [29.2 -<br>34.1] | 51.6 | [48.5 -<br>54.6] | 1.78 |

Finally, GBDP tree reconstruction recognizes both strains as the same species cluster. Besides, the count of features like genomics size and protein count is similar (Figure 1). These observations confirm from the genomic point of view, a previous annotation made by (7,13), in which *B. ciceri* and *B. intermedia* could not be discriminated as different species based solely on traits such as 16S ribosomal gene identity, habitat or lifestyle.

**Figure 1.**
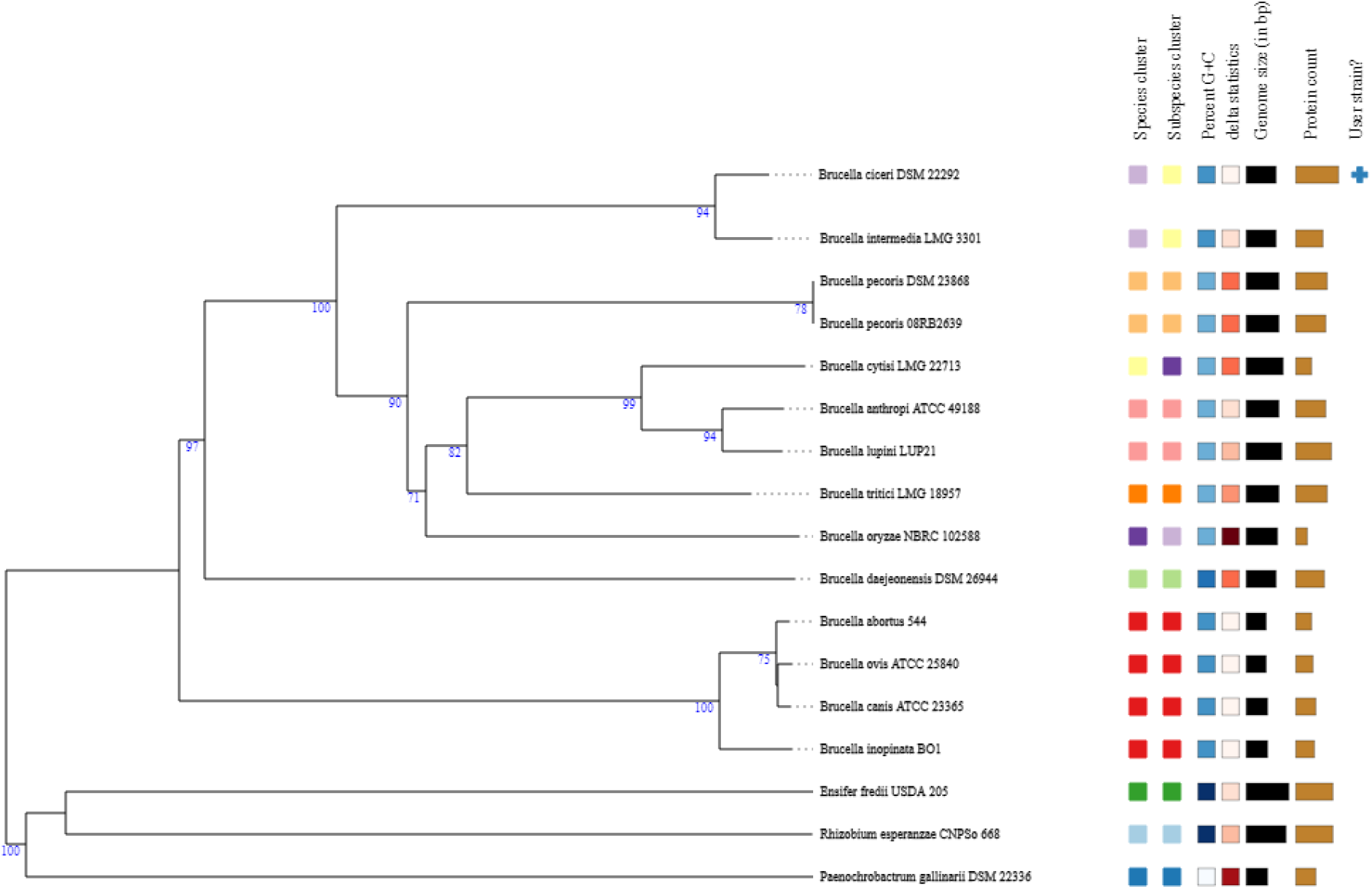
Tree inferred with FastME 2.1.6.1 from GBDP distances calculated from genome sequences. The branch lengths are scaled in terms of GBDP distance formula d5. The numbers above branches are GBDP pseudo-bootstrap support values > 60 % from 100 replications, with average branch support of 80.5 %. The tree was rooted at the midpoint.

In fact, the *iterum O. intermedium*/*O. ciceri* strains represent a diversified clonal population with the capacity to colonize different niches, from pathogenic lifestyle to plant and animal commensal. All these data are highly consistent with that *B. ciceri* DSM 22292^T^ should be reclassified as later heterotypic synonyms of *B. intermedia*. On the other hand, this reclassification does not affect the current clinical terminology, given that *Ochrobactrum intermedium* synonym (correct name *Brucella intermedia*), has been related with major relevance as an opportunistic pathogen.

### Emended species description of *Brucella intermedia*

To the description reported by Velasco et al., (1998), Imran et al., (2010) data are added. Variable reaction to glycogen, Tween 80, raffinose, L-rhamnose, trehalose, a-ketobutyric acid, succinamic acid, glucuronamide, alaninamide, L-threonine, and uridine. A positive reaction for gelatin hydrolysis may occur. Resistant to ampicillin, aztreonam, cefixime, and cephradine. The type strain is CCUG 24694 = CIP 105838 = DSM 17986 = IFO 15820 = LMG 3301 = NBRC 15820= NCTC 12171 = CCUG 57879= DSM 22292.

## Abbreviations

ANI: average nucleotide identity
dDDH: digital DNA–DNA hybridization
Mash D: mutation distance estimated by Mash

